# Increased frequency of recurrent in-frame deletions in new expanding lineages of SARS CoV-2 reflects immune selective pressure

**DOI:** 10.1101/2021.07.04.451027

**Authors:** Arghavan Alisoltani, Lukasz Jaroszewski, Mallika Iyer, Arash Iranzadeh, Adam Godzik

## Abstract

Most of the attention in the surveillance of evolution of SARS-CoV-2 has been centered on single nucleotide substitutions in the spike glycoprotein. We show that in-frame deletions (IFDs) also play a significant role in the evolution of viral genome. The percentage of genomes and lineages with IFDs is growing rapidly and they co-occur independently in multiple lineages, including emerging variants of concerns. IFDs distribution is correlated with spike mutations associated with immune escape and concentrated in proteins involved in interactions with the host immune system. Structural analysis suggests that IFDs remodel viral proteins’ surfaces at common epitopes and interaction interfaces, affecting the virus’ interactions with the immune system. We hypothesize that the increased frequency of IFDs is an adaptive response to elevated global population immunity.

**Summary:** Monitoring of SARS-CoV-2 genome evolution uncovers increased frequency and non-random distribution of in-frame deletions in recently emerged lineages.

## Main text

Deletions, or more generally insertions/deletions (indels), are the second most common modifications in the evolution of viral genomes after single nucleotide polymorphisms (SNPs), and yet receive little attention (*1*). One of the reasons for that is that their consequences on protein structure and function are more challenging to determine than that of single point mutations. Long, loss-of-function deletions removing entire proteins or functional domains could be deleterious (*2*) or attenuating (*3*), however, the effects of shorter, function modifying deletions are mostly unknown. They tend to happen in the loops between secondary structure elements, rarely affecting the overall structure of proteins, but may be altering the binding specificity or protein-protein interaction surfaces (*4*), in few studied examples leading to increased drug resistance and immune escape in viruses (*1, 5*). Their evolutionary dynamics and overall consequences for fitness for any virus, including SARS-CoV-2, remain mostly unaddressed.

Severe acute respiratory syndrome coronavirus 2 (SARS-CoV-2) first emerged in Wuhan, China and subsequently spread worldwide. Its high mutability (*6*), typical for RNA viruses (*7*) but exacerbated by the scale of the COVID-19 pandemic, has resulted in the emergence of multiple lineages. Higher infectiveness and lower efficacy of the current vaccines have been reported for at least two lineages, B.1.351 (*8*) and P.1 (*9, 10*). New lineages combining these two features are still emerging, such as B.1.617.2 now expanding around the world (*11*). Tracking and analysis of emerging new lineages with modified disease phenotypes, dubbed variants of concern (VOCs) (*12*), is crucial for determining the strategies of fighting the COVID-19 pandemic.

Several specific deletions in SARS-CoV-2, such as deletions in the envelope protein (*13*), non-structural protein 1 (NSP1) (*14*), spike glycoprotein(*15*) and accessory ORFs (*16*), have been studied in detail. The NSP1 Δ79-89 was shown to be associated with lower IFN-β levels and non-severe phenotypes (*14*). Regions with recurrent IFDs, called recurrent deletion regions (RDRs), in the N-terminal domain (NTD) of the spike glycoprotein were shown to play a role in immune escape (*15*). These deletions provide an example of a new paradigm of the effects of deletions on viral genomes and proteins – instead of loss-of-function they modify it by remodeling protein surfaces, affecting major antibody epitopes (*17*) and, possibly, protein-protein interaction networks. Our analysis presented here expands on these examples and provides an overview of the dynamics of deletions in the evolution of the SARS-CoV-2 genome.

Large-scale analysis of SARS-CoV-2 genomes (see the Methods section) shows that percentage of genomes with at least one IFD and the fraction of lineages with 1-5 IFDs is growing in time (**Figs. 1A and B**). The recent increase in the number of IFDs was observed in distinct branches of the phylogenetic tree (**Fig. 1C)**, including emerging VOCs. For instance, the B.1.1.7 lineage is defined by 17 founder genome modifications, including three IFDs (NSP6 Δ106-108, spike Δ69-70 and spike Δ144). Additional IFDs and their combinations are found in B.1.1.7 sub-lineages (**Fig. S1**).

**Fig. 1.**
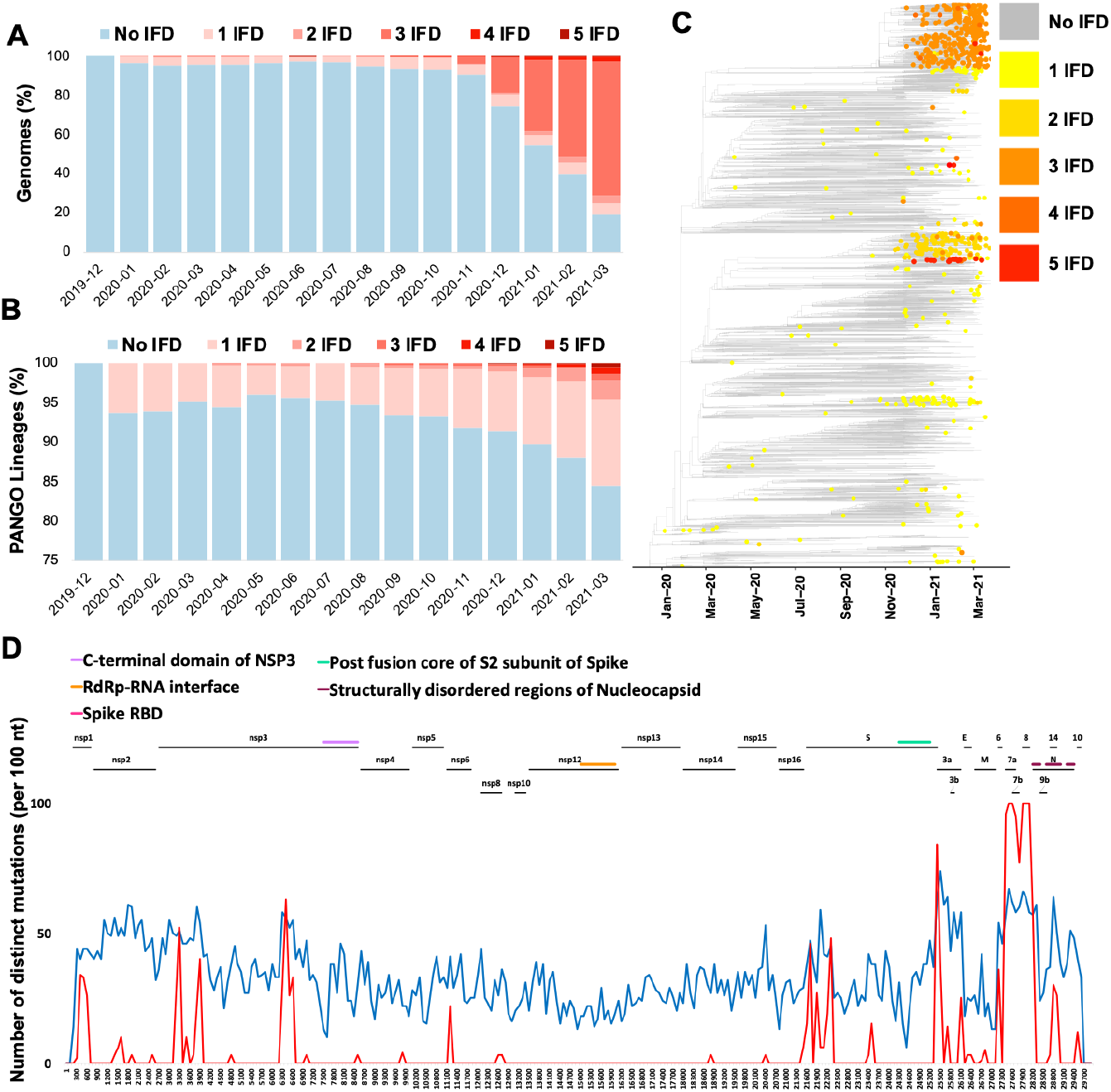
In-frame deletions in SARS-CoV-2 genomes. (A) Percentage of total identified genomes with and without in-frame deletions (IFDs) over time. (B) Percentage of PANGO lineages with and without IFDs over time. (C) Increase in the median number of IFDs in newly emerged lineages illustrated on Nextstrain’s time-resolved phylogenetic tree (D) Distribution of the most common IFDs along the SARS-CoV-2 genome (red) compared to missense SNPs (blue).

Most IFDs are concentrated in specific regions of NSP1, NSP3, NSP6, ORF3a, ORF6, ORF7a, ORF7b, ORF8, nucleocapsid and spike glycoprotein (**Fig. 1D** and **Table S1**), all of which are involved in interactions with the host immune system (*18*). At the same time, proteins involved in the replication–transcription complex show very few or no IFDs (**Fig. 1D** and **Table S1**). Many IFD-prone regions such as the loops in the spike NTD overlap with mutation hotspots (**Fig. 1D**) that are thought to be driven by host immune system pressure (*15, 19, 20*). Therefore, we hypothesize that the emergence of IFDs in the same hotspots is a response to the same pressure. This is supported by the recent studies where both spike-NTD substitutions and indels were demonstrated to accelerate virus adaptation to the host and immune escape (*15, 19, 20*).

Aggregation and recurrence of IFDs in specific regions (Recurrent Deletion Regions or RDRs) of SARS-CoV-2 genomes is determined by an interplay of the protein structural constraints and functional role of specific regions. Most of the RDRs are found on or adjacent to loops forming antibody epitopes (**Fig. 2 and S2**). For instance, NSP6-RDR falls on a short loop between two transmembrane helices that is also a predicted B-cell epitope (**Fig. 2C**). Similarly, NSP1-RDR1, spike RDRs and ORF3a-RDR1, as well as RDRs in other proteins are near or in the loops close to the epitope regions (**Figs. 2 and S2**). Modeling and emerging experimental evidence (*17*) shows such deletions can remodel epitope surfaces, leading to immune escape.

**Fig. 2.**
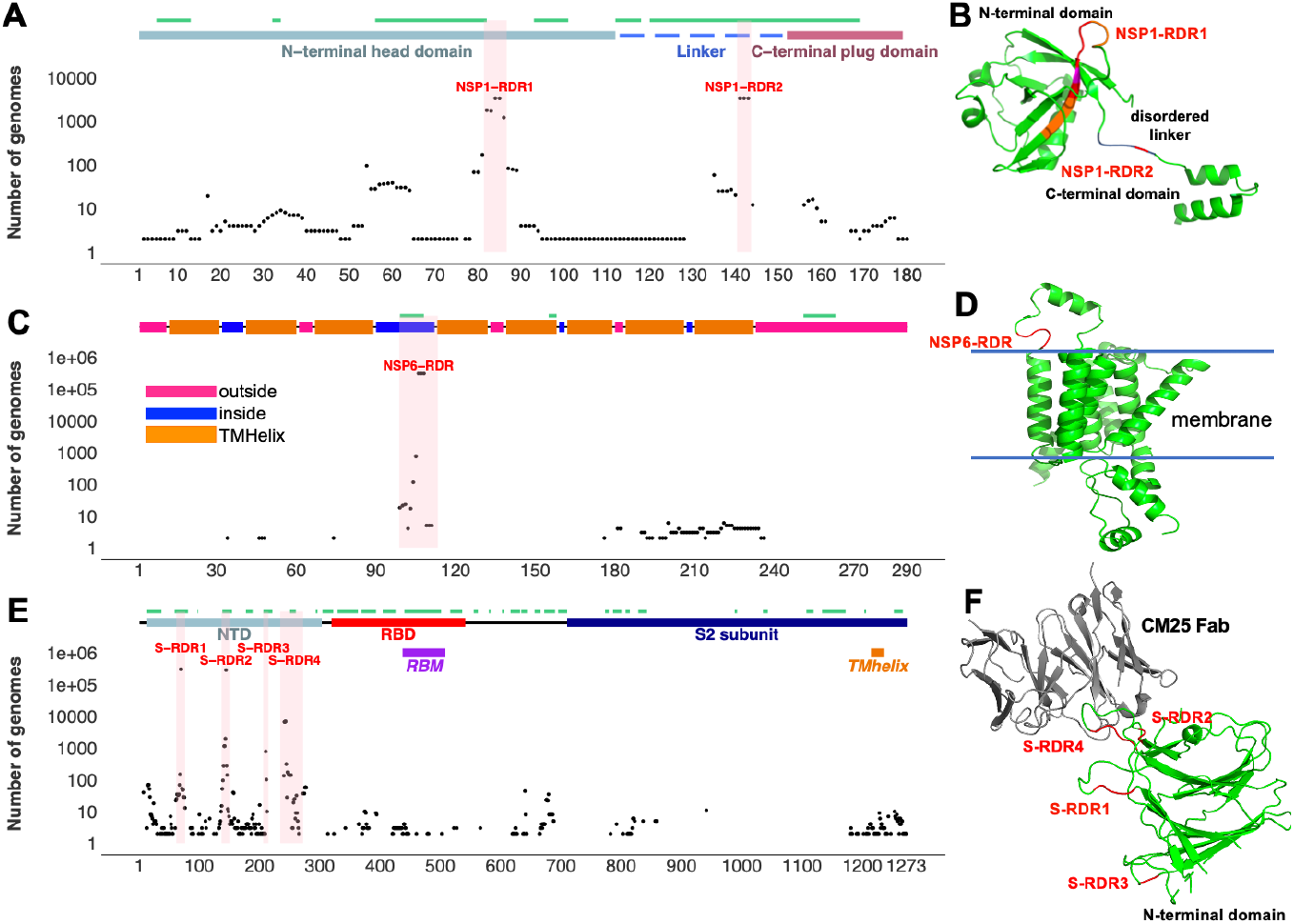
Top SARS-CoV-2 RDRs in the context of protein 3D structure. (A) Distribution of in-frame deletions (IFDs) in NSP1 (B) NSP1-RDRs on protein 3D structure. (C) Distribution of IFDs in NSP6 (D) NSP6-RDR on protein 3D structure. (E) Distribution of IFDs in spike glycoprotein (F) RDRs on the protein 3D structure of the spike glycoprotein N-terminal domain bound to human Fab CM25. IFDs and epitopes are represented as black dots and green lines, respectively. **Table S4** provides details of structures/models used in the figure.

This parallels findings in HIV-1 where deletions in spike glycoprotein regions encoding surface-exposed disordered loops were found to mediate escape from neutralizing antibodies elicited by earlier variants of the virus (*1, 5*).

IFDs positioned on NSP1 and NSP6 RDRs were observed independently in several lineages (**Fig. 3A**) on the SARS-CoV-2 phylogenetic tree (*21*). In the following, we will discuss in detail the RDRs in NSP1, NSP6 and NTD of spike protein. RDRs in other proteins are discussed in **Supplementary Note 1**.

**Fig. 3.**
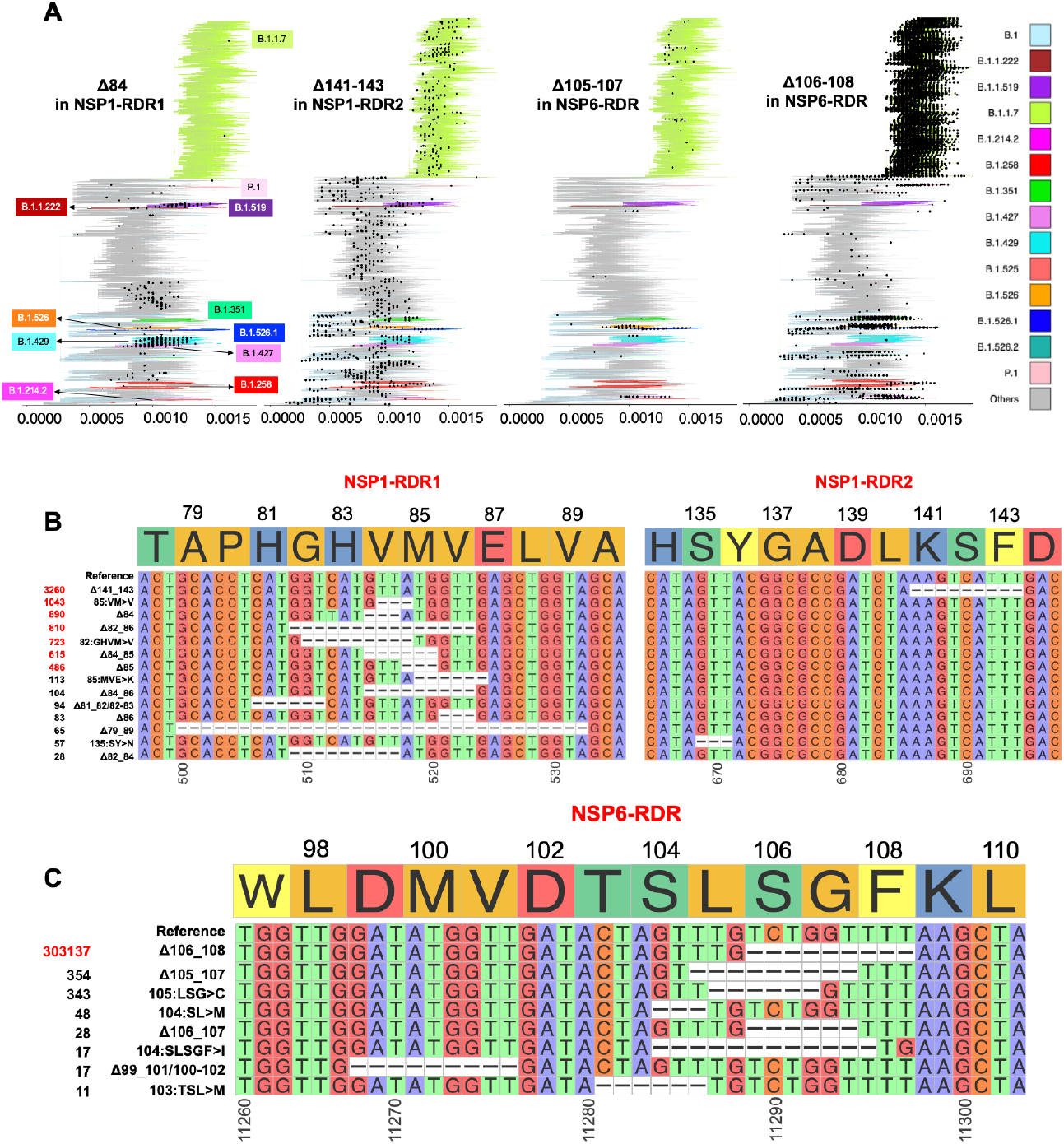
Recurrent deletion regions (RDRs) of NSP1 and NSP6. (A) Global phylogenetic trees of 569,352 SARS-CoV-2 genomes (GISAID as of April 15^th^, 2021) showing recurrence of the most frequent in-frame deletions (IFDs) positioned on NSP1-RDRs and NSP6-RDR as black dots. (B) and (C) represent coordinates of RDRs of NSP1 and NSP6, respectively. The number of genomes containing a specific IFD is provided on the left side of each plot.

Two groups of mutually exclusive NSP1 RDRs (e.g., IFDs Δ84 in NSP1-RDR1 and Δ141-143 in NSP1-RDR2) emerged independently in several lineages such as B.1.1.7, B.1.351, B.1.427, B.1.526 and B.1.258 (**Figs. 3A and 4**). A long version of the IFD in NSP1-RDR1, (Δ79-89) was studied before (*14*), but our analysis indicates that shorter IFDs in this region are recurring more frequently (**Fig. 3A)**. The NSP6 RDR (residues 99-108) is the second most common RDR in SARS-CoV-2, with the Δ106-108 observed in more than 300K genomes (**Fig. 3C**). It independently occurred as a founder modification for several well-known VOCs – B.1.1.7, B.1.351 and P.1, but also some of the newly emerged ones such as B.1.525 in Nigeria and Europe and B.1.526 in New York and Europe (**Figs. 3A and 4**). Signatures of positive selection for NSP6 Δ106-108 were recently reported (*22*). The well-studied IFDs in spike NTD were classified as belonging to RDR1 (residues 60-75), RDR2 (residues 139-146), RDR3 (residues 210-213), and RDR4 (residues 242-248) (*15*). IFDs in NTD-RDR1 and RDR2 are more frequent (compared to RDR3 and RDR4. Several lineages with new spike IFDs (expanding spike-RDR4) and IFDs in other proteins are now emerging (**Fig. S3**).

We observe an increasing number of genomes with independent co-occurrence of multiple spike-IFDs with IFDs in other proteins, especially NSP6-IFDs (**Fig. 4 and Table S3**). NSP6-IFDs independently co-occurred with spike-IFDs located in RDR1 and RDR2 in B.1.1.7 and B.1.525 variants, with IFDs located in RDR2 in B.1.526.1 and B.1.1.318 variants and IFDs in RDR4 in B.1.351 variant (**Fig. 4B and C**). This suggests that different IFD combinations in NSP6 and spike-RDRs might play a role in higher transmissibility, prolonged infection, and immune escape in recent VOCs (**Fig. 4B and C and Table S3)**.

**Fig. 4.**
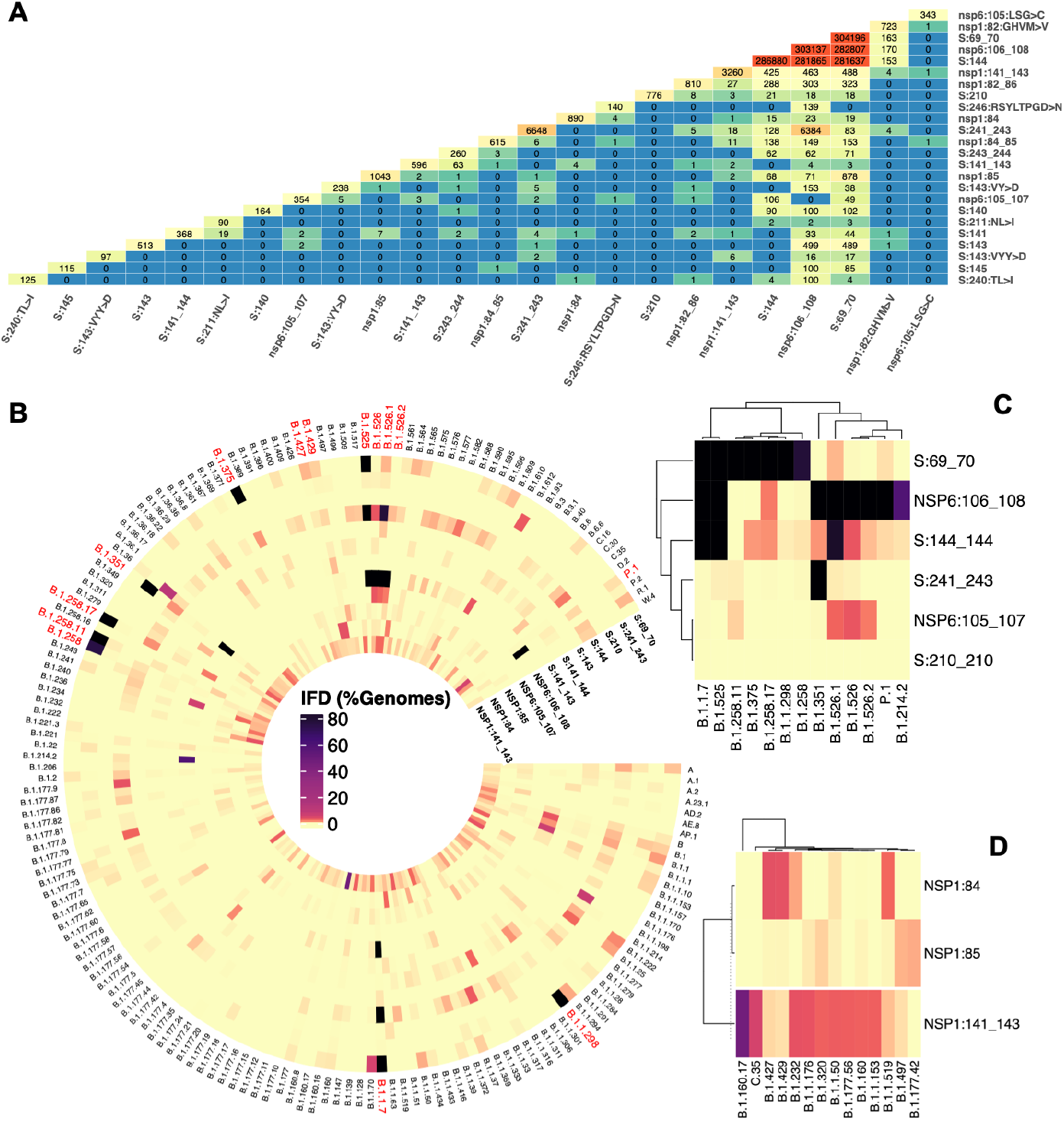
Co-occurrence of in-frame deletions in SARS-CoV-2. **(A)** Co-occurrence of top frequent in-frame deletions (IFDs). (**B**) Independent co-occurrence of top NSP1, NSP6 and spike glycoprotein (S) IFDs in PANGO lineages. The outermost circle shows the name of PANGO lineages with red font indicating lineages harboring recurrent S mutations. The inner circles depict the frequency of IFDs in each lineage. The colors in the inner circles show the % of genomes with a certain IFD with each circle corresponding to one IFD (e.g., black color indicates the IFD is identified in more than 80% of genomes assigned to a given lineage). (**C**) Clustering of NSP6 and S IFDs in most frequent lineages (**D**) Clustering of NSP1-IFDs in most frequent lineages. Data for these heatmaps is provided in **Table S3a-i** which includes additional combinations of IFDs in all lineages.

Independent co-occurrence of IFDs in different lineages might reflect signatures of adaptive evolution by recurrence or recombination. Several VOCs such as B.1.1.7 and B.1.351 which have simultaneous spike and NSP6-IFDs were found to have higher transmissibility, infectivity or immune escape properties than the previously dominant lineages such as B.1.177 (*23*) with almost no IFD (**Fig. 4** and **Table S3**) which could again highlight possible role of simultaneous spike and NSP6-IFDs in immune evasion. Increasing number of IFDs also results in SARS-CoV-2 genome size decrease over time, especially in the recent VOCs (**Fig. S4A** and **B**). Although direct association of genome size with viral fitness is difficult to prove, there is evidence of replicative advantage associated with smaller genome size in RNA viruses (*2, 24, 25*).

In conclusion, we report the increasing rate of recurrent IFDs during the progress of the COVID-19 pandemic, mostly recurring in SARS-CoV-2 proteins involved in interactions with the host immune system, reflecting the host immune selective pressure. Most IFDs are found in recurrent deletion regions, that typically are found in loops close to the epitope regions. Deletions in such regions facilitates immune escape by remodeling the epitope surfaces and prolong infection of these lineages. Such RDRs should be the subject of mutation surveillance as much as common escape mutations. It is likely that increase in the number of IFDs and RDRs in recent lineages is a sign of the virus adapting to the increasing pool of resistant hosts, but more research is needed to decisively prove this point.

## Methods

### Data collection

Multiple sequence alignment (MSA) data, and metadata of complete SARS-CoV-2 genomes (1,028,386) were retrieved from GISAID (https://www.gisaid.org/) as of April 12^th^, 2021. Briefly, full alignment (msa_0412.fasta) provided by GISAID was based on 1,028,386 submissions to GISAID EpiCoV. GISAID pipeline excludes duplicate, low-quality sequences (>5% N content) and incomplete sequences (length <29,000 bp). Then, GISAID pipeline used this cleaned data to create MSA file of 961,734 sequences using MAFFT (*26*) with hCoV-19/Wuhan/WIV04/2019 (EPI_ISL_402124; GenBank: MN996527) used as reference (*27*).

### Identification of in-frame deletions

We used an in-house Perl script to identify variations in each genome based on GISAID MSA file as of April 12^th^, 2021. On top of GISAID’s cutoffs for excluding genomes with high N content and low-quality genomes, we applied additional filtering after variant calling to avoid spurious IFDs and IFDs with shifted positions arising from high N content. Thus, genomes with N content more than 0.05% and more than 200 mutations were excluded resulting in a total of 958,696 SARS-CoV-2 genomes which were used in this study. Additionally, to avoid reporting spurious IFDs arising from sequencing errors or errors in MSA, we used GISAID MSA file with no gaps in reference (obtained with *keep reference length* option) (*26*) to confirm the exact positions of the identified deletions.

### Analyzing SARS-CoV-2 in-frame deletions in the context of PANGO lineages

We used assigned PANGO lineages (*28*) and GISAID (*29*) global divergent tree which includes 569,352 genomes as of April 15^th^, 2021 to investigate the distribution of IFDs across SARS-CoV-2 genomes. We used ggtree R package (*30*) to visualize distributions of IFDs on the GISAID global divergent tree.

### Assessing differences in the rate of in-frame deletions between SARS-CoV-2 proteins

The method we recently used in assessing the significant under-mutated and over-mutated proteins during SARS-CoV-2 evolution (*31*) was used here to identify proteins with a high rate of IFDs. Briefly, we counted the total number of IFDs (except singletons which are usually regarded as unreliable) for each protein (except for NSP11, ORF3b, ORF9b and ORF14 as these are too short for significance analysis). We then used a two-sided binomial test to compare the rate of deletions in each protein to the rate of deletions in the background (all proteins) to identify proteins with high rates of IFDs. Since in our previous study (*31*) we showed that ORF1ab is less frequently mutated and is likely under more stringent purifying selection compared to the genes coding for structural and accessory proteins (ORFs 2-10) we applied an additional statistical comparison of IFD rates to only non-structural proteins to identify NSPs (NSP1-NSP16) with higher rate of IFDs compared to others. For this specific comparison, we run a two-sided binomial test using only ORF1ab (corresponding to proteins NSP1-NSP16) as background. Adjusted p-values (q-values) were calculated using false discovery rate (FDR) method. Proteins with odds ratio more than 1 and q-values less than 0.05 were considered as proteins with significantly increased rates of IFDs.

### Visualization of in-frame deletions on proteins’ 3-dimensional (3D) structures

We used PyMol (*32*), and Coronavirus3D (*33*) for studying and visualization of IFDs in the context of protein 3-dimensional structure (3D). Their 3D coordinates were downloaded from the Protein Data Bank (PDB) (*34*). For proteins with no available 3D structures we used, if available, models predicted by homology modeling or *ab initio* predictions (https://deepmind.com/research/open-source/computational-predictions-of-protein-structures-associated-with-COVID-19, https://zhanglab.dcmb.med.umich.edu/COVID-19/) or SwissModel (*35*), noting in the discussion their hypothetical nature. It should be noted that even for some proteins with available 3D structures we used models predicted by homology modeling or ab initio predictions when the IFDs were located in the regions of the protein with unresolved structures. Information on protein domain boundaries was based on PDB (https://www.rcsb.org/) structures when available or on UniProt and the literature (**Table S4**).

The positions of transmembrane helices for proteins with no available 3D structures were identified with the TMHMM16 server (*36*). IEDB server (Bepipred Linear Epitope Prediction 2.0) (*37*) was used to predict B-cell epitopes for NSP1, NSP2, NSP3, NSP6, Spike, Nucleocapsid, ORF3a, ORF7a, and ORF8 proteins as proteins with significantly increased rate of IFDs.

### Visualization of in-frame deletions (IFDs) on the phylogenetic tree

We mapped the number of IFDs for each genome (1 IFD to maximum 5 IFDs per genome was recorded) on the Nextstrain time-resolved tree (*38*) which includes 3899 genomes sampled between Dec 2019 and March 31^st^, 2021. We used ggtree R package (*30*) to visualize the tree.

### Visualization of in-frame deletions on alignment file

We extracted one representative genome for each of the most frequent IFDs (seen in multiple genomes) positioned on protein RDRs from the GISAID MSA file with no gaps in the reference using an in-house Python script and visualized it using R packages ggmsa and Biostrings and counted the number of genomes harboring each type of IFD.

### Statistical analysis of the most recurrent in-frame deletions in SARS-CoV-2

To identify the most frequent recurrent IFDs in SARS-CoV-2 genomes, we used the GISAID global tree (569,352 genomes as of April 15^th^, 2021) and screened it to find independent occurrences of each IFD (*29*). The most frequent IFDs (observed in at least 400 genomes in all the analyzed lineages) that occurred independently in more than three independent branches of the GISAID global tree and in at least two different lineages were considered recurrent IFDs. Regions with different recurrent IFDs which occurred in adjacent residues (up to 5 residues apart) were called recurrent deletion regions (RDRs). Every RDR involves 2-15 amino acid residues.

We also calculated the recurrence of each IFD as the function of time of sample collection, geographical location, PANGO lineages, and GISAID clades. Briefly, we classified genomes into 16-time bins based on the month and year of the data collection. Similarly, we classified genomes into 6 geographical locations (continents). Genomes were also grouped based on the GISAID clades into 9 groups (clades S, V, L, G, GH, GR, GV, GRY and O), and 1255 different PANGO lineages. Each IFD was counted if it was presented in more than 5 genomes in the given group (the cutoff of five was used to reduce the amount of noise due to deletions with very low counts). We used Circos R package to draw the heatmap of recurrence/co-occurrence of top IFDs in PANGO lineage arranges in the order that approximately reflects the evolutionary history of SARS-CoV-2 lineages.

## Acknowledgments

We gratefully acknowledge the all the authors from the originating laboratories responsible for obtaining the SARS-CoV-2 specimens and the submitting laboratories where genetic sequence data were generated and shared via the GISAID Initiative, on which this research is based. We also acknowledge individual structural biology laboratories that deposited structures used in this work to the PDB. This work is sponsored in part by NIH institutes: NIAID under contract no. HHSN272201700060C and NIGMS by a grant GM118187.

