## supplemental figures and tables for "Increased frequency of recurrent in-frame deletions in new expanding lineages of SARS CoV-2 reflects immune selective pressure"

**This PDF file includes:**

Supplementary Text
Figs. S1 to S5
Tables S1, S2, and S4
References

### Supplementary Text

#### Note 1: Recurrent deletion regions (RDRs) are detected in several proteins

In addition to SARS-CoV-2 NSP1, NSP6 and spike protein RDRs, we identified RDRs and other hotspots for in-frame deletions (IFDs) that are potentially involved in viral-host immune interactions in NSP2, NSP3, ORF3a, ORF7a and ORF8 as well as nucleocapsid protein (**Fig. S5**).

##### *Recurrent IFDs in SARS-CoV-2 non-structural proteins 2 and 3*

Multiple IFDs in the NSP2 RDR (residues 265-270) were observed independently in several lineages (**Fig. S5)**. The NSP2 Δ265-266 is the founder modification of the B.1.573, B.1.1.191, and AN.1 PANGO lineages (**Table S3i**), primarily seen in Canada and Denmark samples. The NSP2 Δ268 is mainly occurring in viral genomes collected from England, Scotland, Northern Ireland and the Netherlands, and it is also the signature mutation of several lineages (**Table S3i**). Interestingly, the NSP2-IFDs are mutually exclusive with IFDs in other proteins - they only co-occur with spike and NSP6-IFDs in very few genomes assigned to B.1.1.7 lineage (**Table S3**). The NSP2 Δ267-268 mostly appeared during early phase of the pandemic and only small portion of the recently collected genomes are harboring other NSP2-IFDs positioned on NSP2-RDR (**Fig. S5**).

Similar to NSP2-RDR, IFDs in NSP3-RDR (residues 1237-1266) are observed in several lineages. The most common IFDs of NSP3 are mostly observed in L.1 PANGO lineage in Canada (Δ1237-1251) and in B.1.1.298 variant from Denmark (Δ1263) where the latter co-occur with NSP1 85:VM>V and spike Δ69-70.

##### *Recurrent IFDs in SARS-CoV-2 Nucleocapsid and accessory proteins ORF3a, ORF7a and ORF8*

Unlike non-structural proteins and spike, accessory proteins have longer IFDs (ORF3a, ORF7a, and ORF8). The most frequent (seen in multiple genomes) but not recurrent (seen in less than two lineages and three independent branches of the GISAID global tree) was ORF7a Δ5-23 that co-occur with NSP6 and spike IFDs in B.1.1.7 lineage (**Fig. S1**). However, the most recurrent IFDs of ORF7a corresponds to residues 58-63, thus defining the ORF7a-RDR (**Fig. S5**). ORF7a The most recurrent and frequent IFD of ORF3a identified at amino acid position 255 (ORF3a-RDR), despite recurring in several lineages including B.1.1.7, is not a signature mutation for any of lineages or sub-lineages (**Fig. S5**). ORF3a Δ255 co-occurred with NSP6 and spike IFDs in B.1.1.7 lineage. The most recurrent and frequent IFD of ORF8 located in residues 63-66.

The RDR of the nucleocapsid protein correspond to residues 202-214, the most frequent IFD in this region (208AR>G) is a signature mutation of B.1.1.318 and also occurred in some B.1.1.7 genomes (Table S3). It co-occurred with three other IFDs in B.1.1.318, including NSP6 Δ106-108, spike Δ144, and ORF7b 44:TNMKF>Y. According to the Coronavirus3D variant tracker this lineage is among the top growing lineages in several countries such as USA, UK, and France (https://coronavirus3d.org/).

***SARS-CoV-2 recurrent deletion regions positioned in or near protein loop regions***

SARS-CoV-2 RDRs in almost all proteins (except ORF7a) are located in or around loop regions and, at the same time, they are near predicted B-cell epitopes, suggesting the selective adaptation of SARS-CoV-2 recurrent IFDs (**Fig. S2).** The NSP2-RDR is located in short surface helix surrounded by loops, and it also corresponds to B-cell epitope region, so this deletion would like lead to the epitope remolding. NSP2 was shown that could disrupt host signaling and might play key role in SARS-CoV-2 pathogenicity. However, more investigations are required to elucidate the role of NSP2 protein and the impact of recurrent IFDs in the virus immune evasion. NSP3 RDR corresponds to group 2 specific marker domain (G2M) a structurally unresolved region of the protein (**Fig. S2C and D**). Based on the predicted model it is located in the loop region of the protein and IFDs in this region occur near B-cell epitopes (**Fig. S2D**). ORF3a-RDR (residues 256-259) is also located in the structurally unresolved region of the protein but based on the predicted structures it also corresponds to a loop region and at the same time positioned on B-cell epitopes. Similar observations were also made for nucleocapsid and ORF8 RDRs **(Fig S2)**. Loop regions of proteins are usually located in the protein surface and often play role in protein-protein interactions and could play role in SARS-CoV-2-host immune interactions.

**Fig. S1**. (A) Combinations of in-frame deletions in the B.1.1.7 PANGO lineage. (B) The genomes with ORF7a Δ5-23 shown as dots in the global phylogenetic tree of 569,352 SARS-CoV-2 genomes retrieved from GISAID as of April 15th, 2021.

**A**

| Combinations of IFDs in B.1.1.7 PANGO Lineage | Number of Genomes |
|---|---|
| nsp6:Δ106_108, S:Δ69_70, S:Δ144 | 268610 |
| nsp6:Δ106_108, S:69:69_70 | 3658 |
| nsp6:Δ106_108, S:Δ69_70, S:Δ144, 7a:5:5_23 | 3637 |
| S:Δ69_70, S:Δ144 | 2901 |
| nsp6:Δ106_108, S:Δ144 | 1122 |
| nsp6:Δ106_108 | 825 |
| nsp6:Δ106_108, S:Δ69_70, S:Δ144, 7a:Δ59_62 | 536 |
| Other | 6681 |

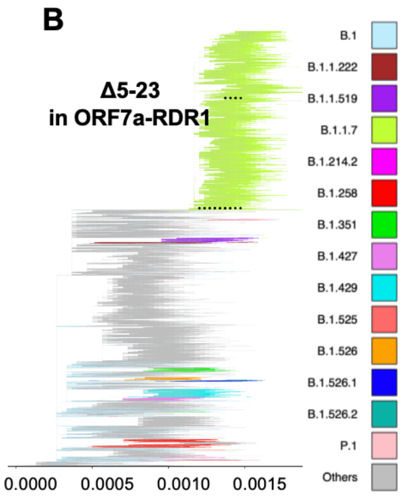

**Fig. S2. SARS-CoV-2 in-frame deletions (IFDs) in the context of protein 3D structure**
(A) Distribution of IFDs in SARS-CoV-2 non-structural protein 2 (NSP2) (B) NSP2 recurrent deletion region (RDR) on protein 3D structure (C) Distribution of IFDs in SARS-CoV-2 NSP3 (D) NSP3-RDR on protein 3D structure (E) Distribution of IFDs in SARS-CoV-2 Nucleocapsid (N) protein (F) N-RDR on protein 3D structure (G) Distribution of IFDs in SARS-CoV-2 ORF3a (H) ORF3a-RDR on protein 3D structure (I) Distribution of IFDs in SARS-CoV-2 ORF7a (J) ORF7-RDR on protein 3D structure. (K) Distribution of deletions in SARS-CoV-2 ORF8 (L) ORF8-RDR on protein 3D structure. IFDs and epitopes are represented as black dots and green lines, respectively. Pink highlighted regions represent RDRs or potential hotspots for recurrent IFDs in each protein. The regions of 3D structure corresponding to RDRs are colored in red. The coordinates of proteins were obtained from different sources (see **Table S4**). Predicted 3D structural models https://zhanglab.ccmb.med.umich.edu/COVID-19/ were used for visualization of recurrent deletion regions in NSP3, ORF3a, and nucleocapsid protein. SP: signal peptide.

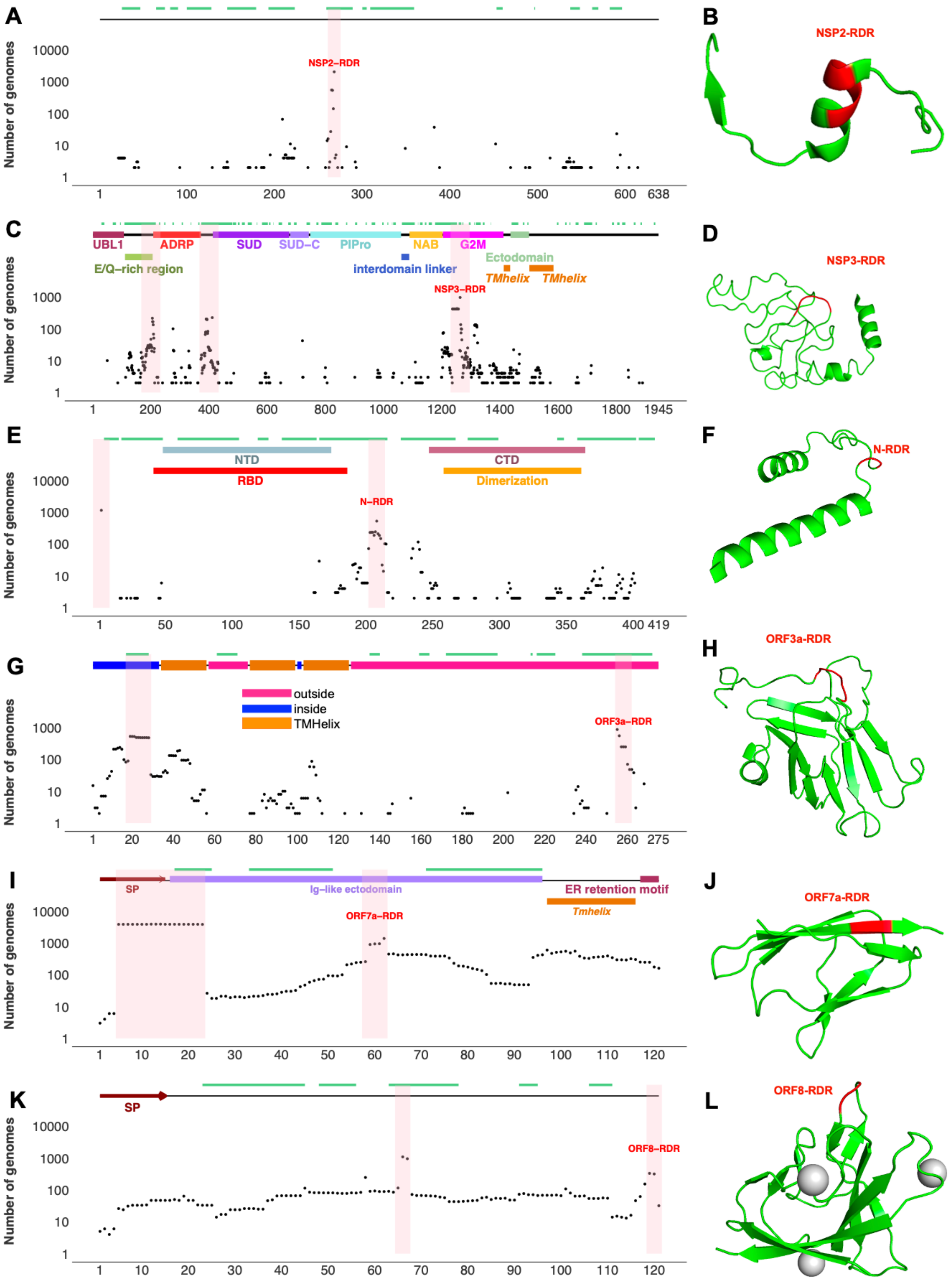
A
B
C
D
E
F
G
H
I
J
K
L
Number of genomes
NSP2-RDR
NSP3-RDR
N-RDR
ORF3a-RDR
ORF7a-RDR
ORF8-RDR
UBL1
ADRP
SUD
SUD-C
PlPro
NAB
G2M
E/Q-rich region
interdomain linker
Ectodomain
TMhelix
NTD
RBD
CTD
Dimerization
outside
inside
TMHelix
SP
Ig-like ectodomain
ER retention motif
Tmhelix

**Fig. S3**. **Recurrent deletion regions (RDRs) and potential hotspots for RDRs in SARS-CoV-2 spike glycoprotein**

**(A)** Combinations of spike IFDs in different genomes **(B)** Potential new RDRs in spike protein **(C)** The most frequent spike IFDs in RDRs1-4 shown as dots in global phylogenetic tree of 569,352 SARS-CoV-2 genomes retrieved from GISAID as of April 15th, 2021.

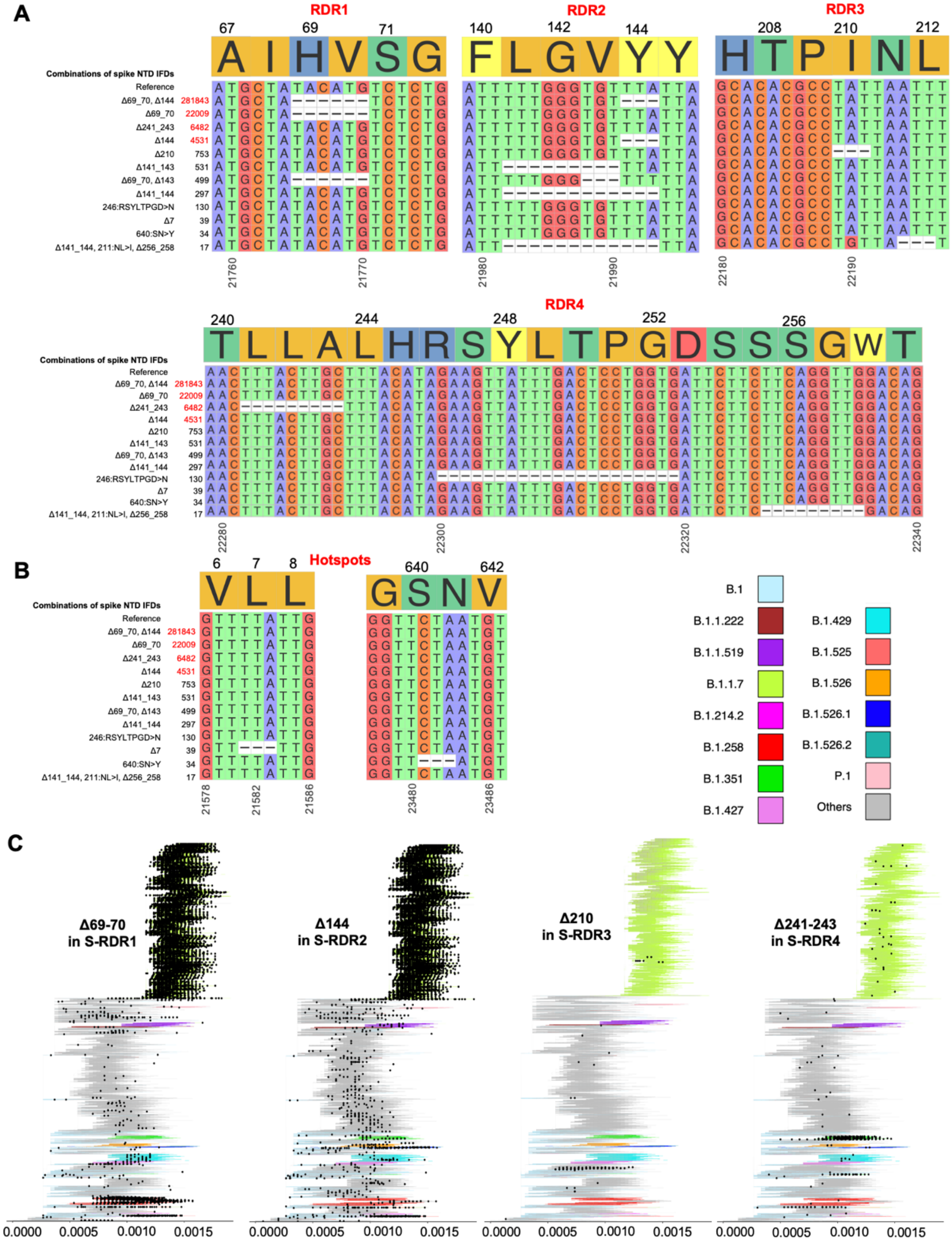

**Fig. S4. SARS-CoV-2 genome shrinkage over the course of pandemic**

(A) Violin plots represent shift in the length of SARS-CoV-2 genomes over the course of the pandemic (B) Difference in the length of SARS-CoV-2 genomes collected before and after the beginning of 2021. (C) Violin plots show top five lineages with the smallest genome length. P-values calculated using Wilcoxon–Mann–Whitney test. (D) Median length of SARS-CoV-2 genomes in the most abundant lineages (>2000 genomes in each represented lineage). nt: nucleotide

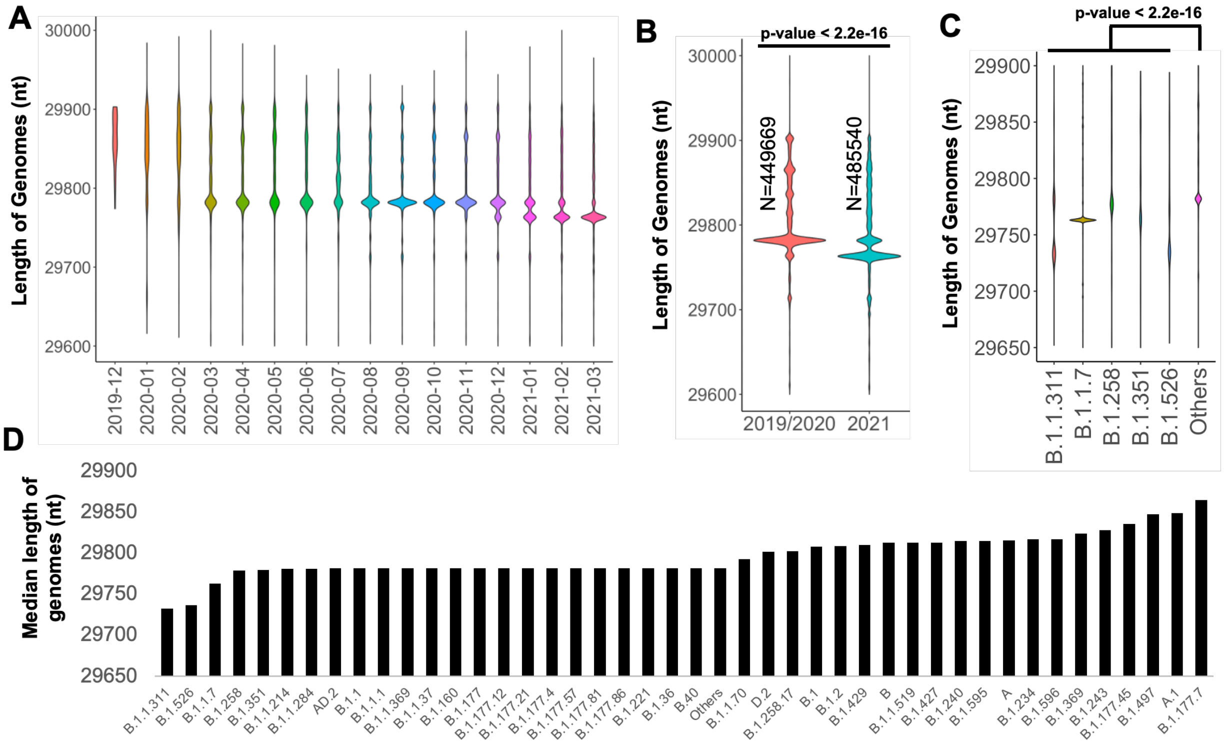

**Fig. S5**. In-frame deletions (IFDs) shown as dots in global phylogenetic tree of 569,352 SARS-CoV-2 genomes retrieved from GISAID as of April 15th, 2021. Only most frequent and recurrent IFDs located in recurrent deletion regions of SARS-CoV-2 proteins are presented.

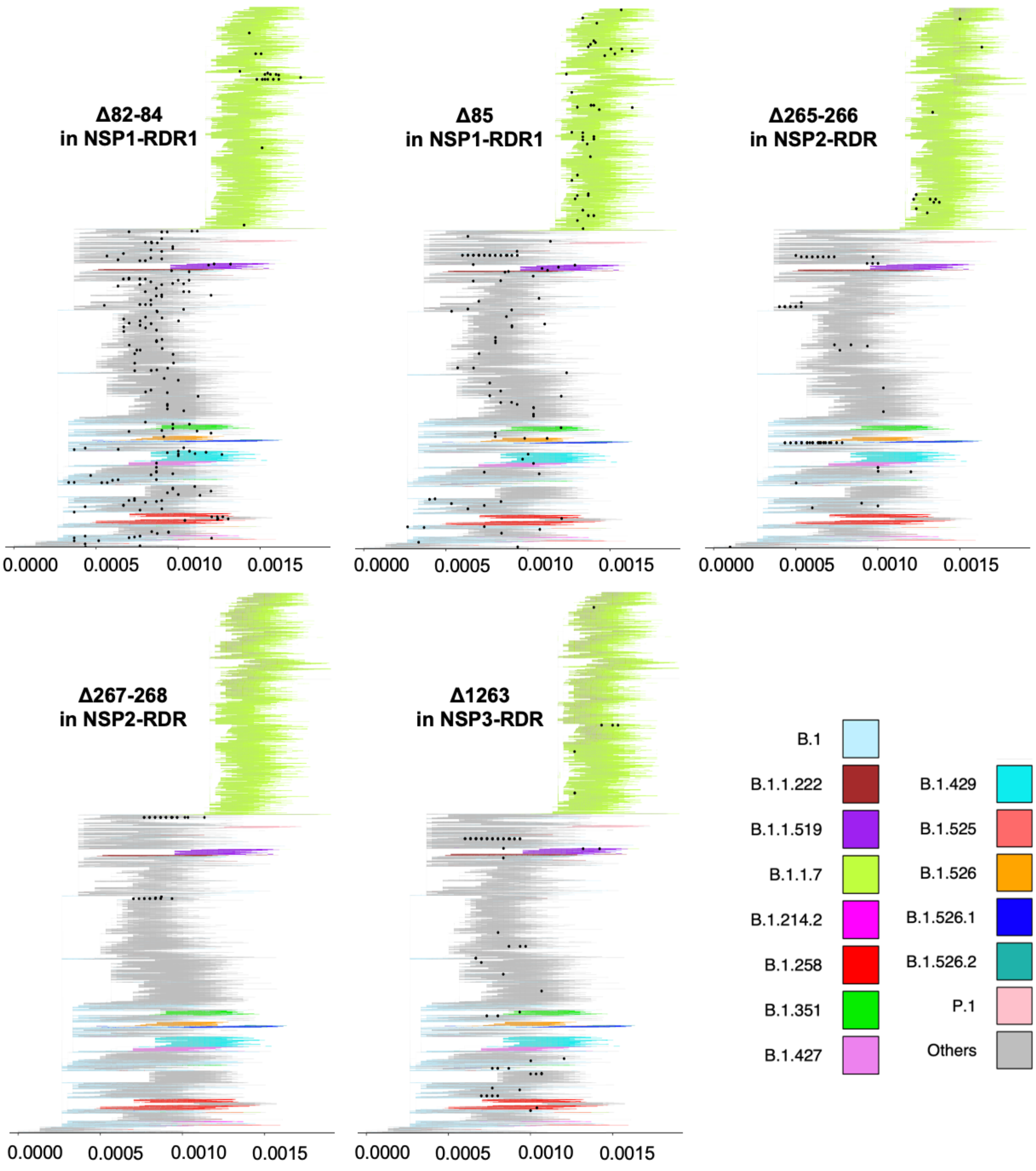

**Fig. S5. Continued…**

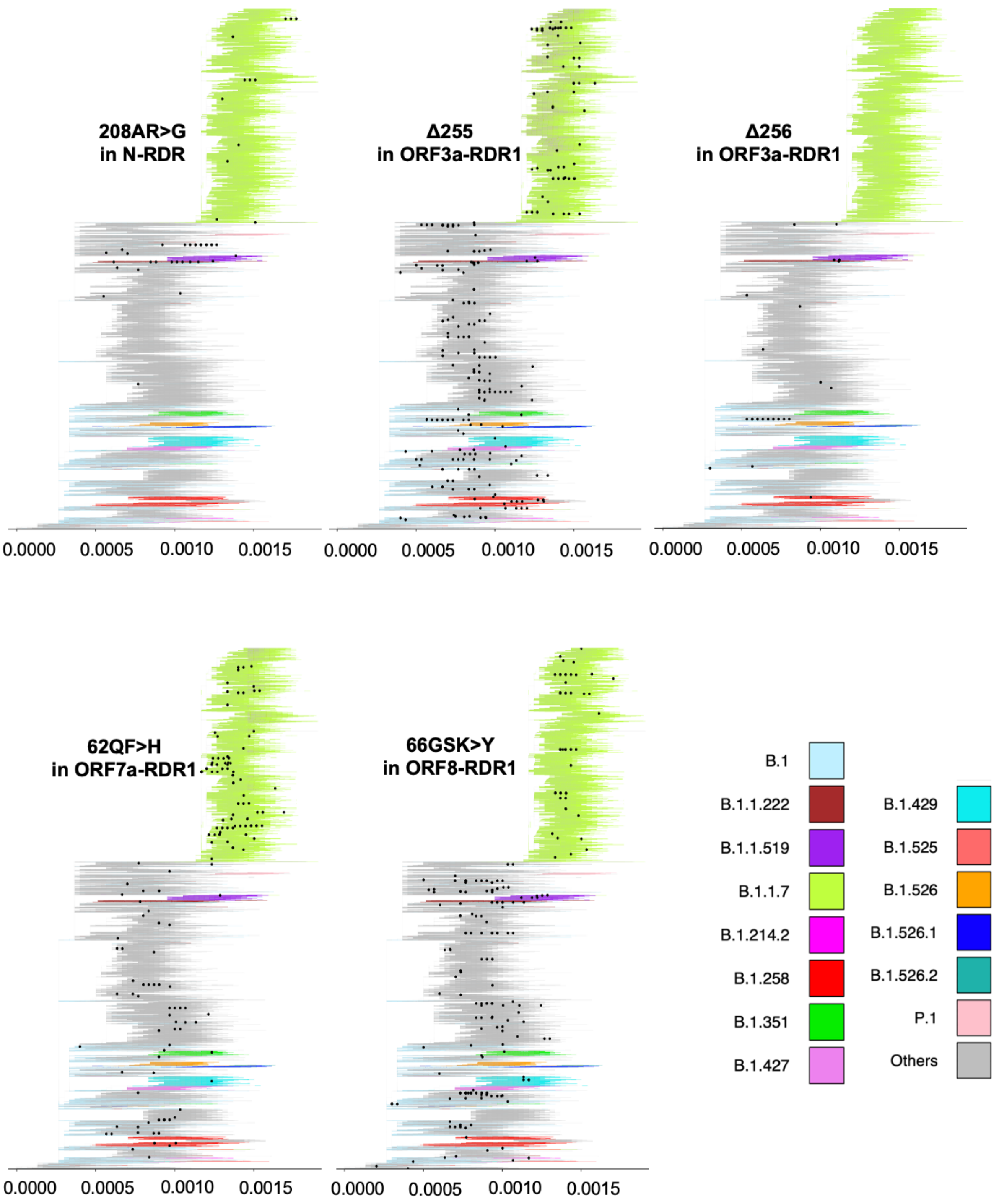

**Table S1.** Comparison of frequencies of in-frame deletions (IFDs) in SARS-CoV-2 proteins using two-sided binomial test (only IFDs observed in at least 2 genomes were included to eliminate spurious mutations). Only PANGO lineages harboring the most frequent IFD (observed in at least 10 genomes of each lineage) are reported. Bold font indicates proteins with significantly increased rate of IFDs (q-value<0.05 and Odds ratio>1).

| Protein | No. IFDs (Singletons excluded) | All proteins as background | | ORF1ab as background | | Most frequent IFD | No. genomes with most frequent IFDs | Top lineages with most frequent IFD in each protein |
|---|---|---|---|---|---|---|---|---|
| | | Odds ratio | q-value (FDR adjusted p-value) | Odds ratio | q-value (FDR adjusted p-value) | | | |
| **NSP1** | **92** | **2.19** | **4.47E-11** | **6.19** | **6.52E-23** | **141_143** | **3260** | **B.1.1.214, B.1.517, AE.1, B.1.1.176, B.1.240, B.1.177.56, B.1.1, B.1.177.21, B.1.595.1, B.1.279, B.1.177, B.1.35, B.1.526, B.1.234, B.1.221, B.1.258, B.1.372, B.1.1.284, B.1.1.519, A.27, B.1.351, B.1.160.17, B.1.1.70, B.1.258.17, B.1.177.4, B.1.429, B.1.1.7, B.1.427, B.1.2, B.1.232, B.1.526.2, B, B.1, B.1.243, B.1.160, D.2, B.1.526.1, B.1.1.372, B.1.195, B.1.1.37, B.1.596, B.1.1.50, C.35** |
| NSP2 | 97 | 0.65 | 6.08E-06 | 1.03 | 9.03E-01 | 268 | 1908 | B.23, B.1.1, B.10, B, B.11, B.47, B.49, B.57, B.1, B.27, B.52, B.45, B.55, B.15 |
| **NSP3** | **410** | **0.90** | **2.38E-02** | **1.72** | **1.34E-12** | **1263** | **963** | B.1.8, B.1.1.7, B.1.234, B.1, B.1.1.298 |
| NSP4 | 65 | 0.56 | 2.92E-07 | 0.59 | 6.27E-02 | 435 | 225 | B.1.1.301, B.1, B.1.2 |
| NSP5 | 16 | 0.22 | 1.20E-14 | 0.07 | 1.01E-04 | 168 | 4 | |
| **NSP6** | **58** | **0.86** | **2.89E-01** | **1.96** | **3.08E-03** | **106-108** | **303137** | **B.1.526, B.1.383, B.1, B.1.1.7, B.1.1, B.1.526.1, B.1.526.2, B.1.351, B.1.525, B.1.1.318, B.1.243.1, B.1.160, B.1.214.2, B.1.1.1, B.1.470, P.1, B.1.2, B.1.221, B.1.241, B.1.258.17, B.1.177, B.1.214** |
| NSP7 | 4 | 0.21 | 7.13E-05 | 0.55 | 6.87E-01 | 45 | 3 | |
| NSP8 | 14 | 0.30 | 8.80E-08 | 0.46 | 1.51E-01 | 179:NL>I | 25 | B.1.1 |
| NSP9 | 12 | 0.46 | 4.05E-03 | 0.81 | 1.00E+00 | 33 | 26 | B.1.1.44 |
| NSP10 | 8 | 0.25 | 9.57E-07 | 0.00 | 6.68E-03 | 101 | 1 | |
| NSP12 | 74 | 0.34 | 1.37E-30 | 0.27 | 5.67E-08 | 643 | 11 | B.1.1.7 |
| NSP13 | 33 | 0.24 | 7.31E-28 | 0.26 | 3.37E-05 | 206 | 9 | |
| NSP14 | 60 | 0.49 | 4.72E-10 | 0.35 | 8.12E-04 | 287 | 46 | B.1.1.7 |
| NSP15 | 45 | 0.56 | 2.41E-05 | 0.46 | 3.78E-02 | 269 | 30 | B.1 |
| NSP16 | 26 | 0.37 | 5.81E-09 | 0.38 | 2.67E-02 | 297 | 27 | |
| **Spike** | **340** | **1.14** | **1.06E-02** | **-** | **-** | **69-70** | **304196** | **B.1.1.37, A.28, B.1.429, B.1.258.5, B.1.1.298, B.1.177** |
| **ORF3a** | **176** | **2.74** | **6.34E-31** | **-** | **-** | **255** | **864** | **B.1.221, B.1.2, B.1.1, B.1.1.7, C.1, B.1.177.21, B.1.258, B.1.1.38, B.1.1.291, B.1.258.17, B.1, B.1.1.375, B.1.177, B.1.517, C.8** |
| E | 20 | 1.14 | 5.69E-01 | - | - | 75 | 46 | B.1.1.7 |
| M | 33 | 0.64 | 7.57E-03 | - | - | 91-98 | 8 | |
| **ORF6** | **43** | **3.02** | **1.32E-09** | **-** | **-** | **2** | **1136** | **B.1.177, B.1.525, B.1** |
| **ORF7a** | **311** | **11.01** | **3.21E-210** | **-** | **-** | **5-23** | **3763** | **B.1.1.7** |
| **ORF7b** | **62** | **6.18** | **5.695E-28** | **-** | **-** | **44:*TNMKF>Y** | **279** | **B.1.1, B.1.1.318** |
| **ORF8** | **126** | **4.46** | **2.77E-41** | **-** | **-** | **GSK>E** | **767** | **B.1, B.1.2, B.1.36.36, B.1.1, B.1.1.413, B.1.1.214, B.1.36.10, B.1.1.7, AD.2, B.1.177** |
| **N** | | **1.44** | **3.89E-05** | **-** | **-** | **2:SD>Y** | **1149** | **B.1.525, B.1, B.1.177** |
| ORF10 | 8 | 0.90 | 1.00E+00 | - | - | 9-10 | 1025 | B.1.128 |

**Table S2.** The most frequent deletions in SARS-CoV-2 genome - the IFDs which occurred in at least three different clades (either PANGO lineages or GISAID clades).

| IFDs | Number of genomes with IFDs | PANGO lineages (total 1255) | GISAID Clades (total 9) | Region (total 6 continents) | Time bins (total 16 months) |
|---|---|---|---|---|---|
| **nsp1:141_143(686_694)** | 3260 | 67 | 9 | 6 | 13 |
| **S:144_144(21991_21993)** | 286880 | 41 | 9 | 6 | 14 |
| **S:69_70(21765_21770)** | 304196 | 42 | 7 | 6 | 13 |
| **nsp1:82_86(508_522)** | 810 | 21 | 7 | 5 | 13 |
| **8b:66:GSK>E(28090_28095)** | 767 | 19 | 5 | 6 | 13 |
| **nsp6:106_108(11288_11296)** | 303137 | 29 | 6 | 6 | 8 |
| **nsp1:84_85(515_520)** | 615 | 17 | 7 | 5 | 13 |
| **S:141_144(21982_21993)** | 368 | 11 | 7 | 5 | 13 |
| **nsp2:268_268(1605_1607)** | 1908 | 16 | 6 | 5 | 8 |
| **nsp1:82:GHVM>V(510_518)** | 723 | 16 | 5 | 5 | 13 |
| **nsp1:85_85(518_520)** | 486 | 14 | 7 | 3 | 13 |
| **3a:255_255(26155_26157)** | 864 | 26 | 5 | 4 | 12 |
| **S:241_243(22281_22289)** | 6648 | 10 | 6 | 5 | 7 |
| **nsp1:85_85(516_518)** | 1043 | 9 | 5 | 4 | 11 |
| **3a:256:VN>D(26159_26161)** | 370 | 9 | 5 | 3 | 8 |
| **nsp4:435_435(9858_9860)** | 225 | 8 | 5 | 3 | 12 |
| **7a:62:QF>H(27579_27581)** | 388 | 7 | 5 | 3 | 9 |
| **nsp3:1263_1263(6506_6508)** | 963 | 6 | 5 | 4 | 8 |
| **7b:13_13(27792_27794)** | 271 | 8 | 6 | 3 | 7 |
| **S:210_210(22189_22191)** | 776 | 6 | 4 | 4 | 9 |
| **S:141_143(21981_21989)** | 596 | 7 | 4 | 4 | 7 |
| **nsp1:84_86(514_522)** | 104 | 5 | 5 | 4 | 7 |
| **S:243_244(22289_22294)** | 260 | 7 | 5 | 3 | 7 |
| **nsp1:84_84(514_516)** | 890 | 15 | 4 | 2 | 8 |
| **nsp2:265_266(1598_1603)** | 517 | 6 | 4 | 2 | 11 |
| **nsp1:85:MVE>K(519_524)** | 113 | 2 | 5 | 3 | 8 |
| **nsp6:105_107(11283_11291)** | 354 | 9 | 4 | 3 | 4 |
| **7b:14_14(27795_27797)** | 127 | 5 | 5 | 2 | 6 |
| **7a:103_103(27698_27700)** | 110 | 2 | 6 | 3 | 5 |
| **S:143:VY>D(21990_21992)** | 238 | 5 | 4 | 3 | 3 |
| **S:143:VYY>D(21990_21995)** | 97 | 5 | 5 | 2 | 6 |
| **7a:97_97(27682_27684)** | 102 | 4 | 4 | 2 | 7 |
| **3a:256_259(26158_26169)** | 144 | 5 | 4 | 2 | 6 |
| **7a:102_103(27695_27700)** | 107 | 4 | 4 | 2 | 6 |
| **nsp1:81_82(505_510)** | 94 | 3 | 4 | 3 | 6 |
| **nsp3:276:NG>R(3546_3548)** | 86 | 4 | 5 | 2 | 6 |
| **8b:58_58(28065_28067)** | 158 | 4 | 4 | 2 | 6 |
| **nsp1:54_54(423_425)** | 77 | 4 | 4 | 3 | 5 |
| **8b:66:GS>A(28090_28092)** | 135 | 4 | 2 | 3 | 6 |
| **nsp2:209:NE>K(1431_1433)** | 61 | 4 | 4 | 3 | 5 |
| **7b:2_2(27759_27761)** | 84 | 3 | 3 | 3 | 7 |
| **8b:66_67(28087_28092)** | 77 | 5 | 3 | 3 | 4 |
| **N:208:AR>G(28896_28898)** | 364 | 4 | 3 | 5 | 4 |
| **S:140_140(21980_21982)** | 164 | 3 | 4 | 3 | 5 |
| **6:22_30(27264_27290)** | 138 | 3 | 4 | 2 | 5 |
| **S:145_146(21993_21998)** | 115 | 4 | 3 | 3 | 4 |
| **nsp2:267_268(1604_1609)** | 129 | 3 | 2 | 3 | 6 |
| **6:2_2(27205_27207)** | 1136 | 3 | 2 | 5 | 4 |
| **nsp3:1265:SL>I(6513_6515)** | 90 | 3 | 4 | 3 | 4 |
| **N:2:SD>Y(28278_28280)** | 1149 | 3 | 1 | 5 | 4 |
| **3a:11_13(25423_25431)** | 119 | 3 | 4 | 2 | 4 |
| **nsp1:135:SY>N(669_671)** | 57 | 3 | 3 | 3 | 3 |
| **N:239_239(28988_28990)** | 60 | 5 | 3 | 2 | 3 |
| **nsp3:205_208(3331_3342)** | 82 | 3 | 2 | 2 | 6 |
| **3a:14_15(25430_25435)** | 88 | 2 | 4 | 2 | 4 |
| **nsp3:1313_1321(6655_6681)** | 82 | 2 | 4 | 2 | 4 |
| **N:209:RMAG>S(28899_28907)** | 84 | 3 | 2 | 2 | 4 |
| **7a:59_62(27568_27579)** | 581 | 1 | 4 | 2 | 4 |
| **nsp1:86_86(521_523)** | 83 | 1 | 5 | 2 | 3 |
| **S:211:NL>I(22194_22196)** | 90 | 2 | 3 | 3 | 3 |
| **3a:19_28(25446_25475)** | 452 | 3 | 1 | 2 | 4 |

**Table S4.** Structures, models and details of protein functional regions used in Figures 2 and S2 paper. Structural models for visualization of RDRs on NSP3, NSP6, ORF3a and nucleocapsid protein 3D structures obtained from https://zhanglab.ccmb.med.umich.edu/COVID-19/. PDB: Protein Data Bank; DB: database

| Protein | Annotation | Start | End | Annotation/Source | PDB IDs |
|---|---|---|---|---|---|
| ORF3a | inside | 57 | 76 | UniProt DB/ PDB/TMHMM server | 6xdcA |
| ORF3a | inside | 126 | 275 | UniProt DB/ PDB/TMHMM server | 6xdcA |
| ORF3a | outside | 1 | 33 | UniProt DB/ PDB/TMHMM server | 6xdcA |
| ORF3a | outside | 100 | 102 | UniProt DB/ PDB/TMHMM server | 6xdcA |
| ORF3a | TMhelix | 34 | 56 | UniProt DB/ PDB/TMHMM server | 6xdcA |
| ORF3a | TMhelix | 77 | 99 | UniProt DB/ PDB/TMHMM server | 6xdcA |
| ORF3a | TMhelix | 103 | 125 | UniProt DB/ PDB/TMHMM server | 6xdcA |
| ORF7a | SP | 1 | 15 | UniProt DB/ PDB/TMHMM server | 6xdcA |
| ORF7a | Ig-like ectodomain | 16 | 96 | UniProt DB/ PDB/(*1*) | 6w37A |
| ORF7a | TMhelix | 97 | 116 | UniProt DB/ PDB/(*1*) | 6w37A |
| ORF7a | typical ER retention motif | 117 | 121 | UniProt DB/ PDB/(*1*) | 6w37A |
| ORF8 | SP | 1 | 15 | UniProt DB/ | 7jx6A |
| nsp1 | nsp1 head, N-terminal domain | 1 | 112 | UniProt DB/ | 7k3nA |
| nsp1 | nsp1 linker | 113 | 151 | UniProt DB | NA |
| nsp1 | plug domain, C-terminal domain | 152 | 179 | UniProt DB/ PDB | 7k5i1 |
| nsp2 | | 1 | 638 | PDB | 7MSW |
| nsp3 | UBL1 | 1 | 107 | UniProt DB/ PDB | 7kagA |
| nsp3 | E/Q-rich region | 111 | 206 | UniProt DB | NA |
| nsp3 | ADRP | 208 | 372 | UniProt DB/ PDB | 6w02A |
| nsp3 | SUD | 413 | 676 | UniProt DB/ PDB | 2w2gA |
| nsp3 | SUD-C | 679 | 743 | UniProt DB/ PDB | 2kafA |
| nsp3 | PIPro | 748 | 1060 | UniProt DB/ PDB | 6w9cA |
| nsp3 | interdomain linker | 1061 | 1088 | UniProt DB | NA |
| nsp3 | NAB | 1089 | 1203 | UniProt DB/ PDB | 2k87A |
| nsp3 | G2M | 1204 | 1412 | UniProt DB | NA |
| nsp3 | Ectodomain | 1436 | 1500 | UniProt DB | NA |
| nsp3 | TMhelix | 1413 | 1435 | UniProt DB | NA |
| nsp3 | TMhelix | 1501 | 1584 | UniProt DB | NA |
| nsp6 | inside | 1 | 11 | TMHMM server | NA |
| nsp6 | inside | 61 | 66 | TMHMM server | NA |
| nsp6 | inside | 133 | 138 | TMHMM server | NA |
| nsp6 | inside | 180 | 183 | TMHMM server | NA |
| nsp6 | inside | 233 | 290 | TMHMM server | NA |
| nsp6 | outside | 32 | 40 | TMHMM server | NA |
| nsp6 | outside | 90 | 112 | TMHMM server | NA |
| nsp6 | outside | 159 | 161 | TMHMM server | NA |
| nsp6 | outside | 207 | 209 | TMHMM server | NA |
| nsp6 | TMhelix | 12 | 31 | TMHMM server | NA |
| nsp6 | TMhelix | 41 | 60 | TMHMM server | NA |
| nsp6 | TMhelix | 67 | 89 | TMHMM server | NA |
| nsp6 | TMhelix | 113 | 132 | TMHMM server | NA |
| nsp6 | TMhelix | 139 | 158 | TMHMM server | NA |
| nsp6 | TMhelix | 162 | 179 | TMHMM server | NA |
| nsp6 | TMhelix | 184 | 206 | TMHMM server | NA |
| nsp6 | TMhelix | 210 | 232 | TMHMM server | NA |
| S | SP | 1 | 12 | UniProt DB/ PDB | 7KNB |
| S | NTD | 13 | 303 | UniProt DB/ PDB | 71qwA/7KNB |
| S | RBD | 319 | 541 | UniProt DB/ PDB | 6lzgB/7KNB |
| S | S2 subunit | 710 | 1274 | UniProt DB/ PDB | 7KNB |
| S | TMhelix | 1214 | 1236 | UniProt DB/ PDB | 7KNB |
| S | RBM | 437 | 508 | UniProt DB/ PDB | 7KNB |
| N | NTD | 48 | 174 | UniProt DB/ PDB | 6m3mA/7KNB |
| N | CTD | 247 | 364 | UniProt DB/ PDB | 6wjiA/7KNB |
| N | RBD | 41 | 186 | UniProt DB/ PDB | 7KNB |
| N | Dimerization | 258 | 361 | UniProt DB/ PDB | 7KNB |
